# Simultaneous broad protection against Ebola Sudan, Marburg and Lassa viruses conferred by a DNA primed MVA-vectored multivalent vaccine

**DOI:** 10.64898/2026.03.18.712579

**Authors:** Darwyn Kobasa, Martina Pfranger, Nina Krause, Sofiya Fedosyuk, Lara Wiegand, Ingo Jordan, Volker Sandig, Christina Leupold, Benedikt Asbach, Daniel Storisteanu, Anja Kalender, Lara Scheer, David Brenner, Jérémie Prévost, Nikesh Tailor, Robert Vendramelli, Bryce Warner, Thang Truong, Srivatsan Parthasarathy, Jonathan Holbrook, George Carnell, Sneha B. Sujit, Sruthika Ashokan, Andrew Chan, Charles Whittaker, Chinedu Ugwu, Christian Happi, Simon Frost, Rebecca Kinsley, David Safronetz, Ralf Wagner, Jonathan L. Heeney

## Abstract

Sub-Saharan Africa continues to experience recurrent outbreaks of zoonotic viral diseases that spill over unpredictably from animal reservoirs into human populations. In many regions, mpox co-circulates with viral hemorrhagic fevers (VHFs) caused by Ebola Sudan virus (SUDV), Marburg virus (MARV), and Lassa fever virus (LASV). Overlapping clinical syndromes that these VHF cause challenge surveillance, diagnostics and timely deployment of effective countermeasures. A single vaccine capable of protecting against these biologically and genetically distinct pathogens would markedly reduce the cost and complexity of outbreak response, lessen dependence on emergency international aid, and strengthen long-term health system resilience. Here, we report on the development of an MVA-based mpox vaccine engineered to express computationally designed, broad-coverage antigens targeting SUDV, MARV and LASV. In preclinical challenge studies, this multivalent vaccine elicited robust immune responses and conferred significant protection against lethal infection from all three pathogens in parallel challenge experiments. These findings establish preclinical proof-of-concept for a single, broadly protective VHF vaccine and support its clinical development for deployment across diverse settings in Sub-Saharan Africa.

**Significance:** Outbreak control in Sub-Saharan Africa is challenged by the co-circulation of different high consequence human infections such as mpox and diverse viral hemorrhagic fevers (VHF) such as SUDV, MARV, and LASV pathogens. These VHFs have overlapping early clinical syndromes, complicating triage and delaying effective targeted interventions. We developed a single MVA-based vaccine encoding computationally designed, conserved antigens from all three VHFs encoded within the MVA vector analogous to the licensed mpox vaccine. In simultaneous challenge models, this multivalent vaccine elicited robust humoral and cellular responses and conferred significant protection against lethal infection by each hemorrhagic fever pathogen. This work provides preclinical proof-of-concept for a unified, broadly protective countermeasure compatible with existing MVA-mpox vaccine manufacturing and deployment experience. By reducing dependence on rapid differential diagnostics and streamlining logistics relative to maintaining multiple pathogen-specific vaccine stockpiles, this approach can lower costs, accelerate response, and increase equity of access during syndromic outbreaks. The platform’s engineered antigen breadth and human safety profile of MVA together support a pragmatic translational pathway toward clinical evaluation and regional readiness for zoonotic spillover events that are intensifying with human and environmental changes.

## Introduction

Emerging and re-emerging zoonotic viral diseases that spill over unpredictably from animal reservoirs continue to impose severe health and economic burdens across Sub-Saharan Africa. Mpox and several viral hemorrhagic fevers (VHFs), including those caused by Sudan virus (SUDV), Marburg virus (MARV), and Lassa fever virus (LASV) often occur in overlapping geographic regions and produce clinically similar diseases early disease presentations. These pathogens are associated with high transmissibility, severe case fatality rates, and the potential for rapid regional spread (Figure 1). Together they place sustained pressure on fragile health systems with limited surge capacity, contributing to increased mortality, disruptions in essential services, and prolonged societal and economic instability.

**Figure 1.**
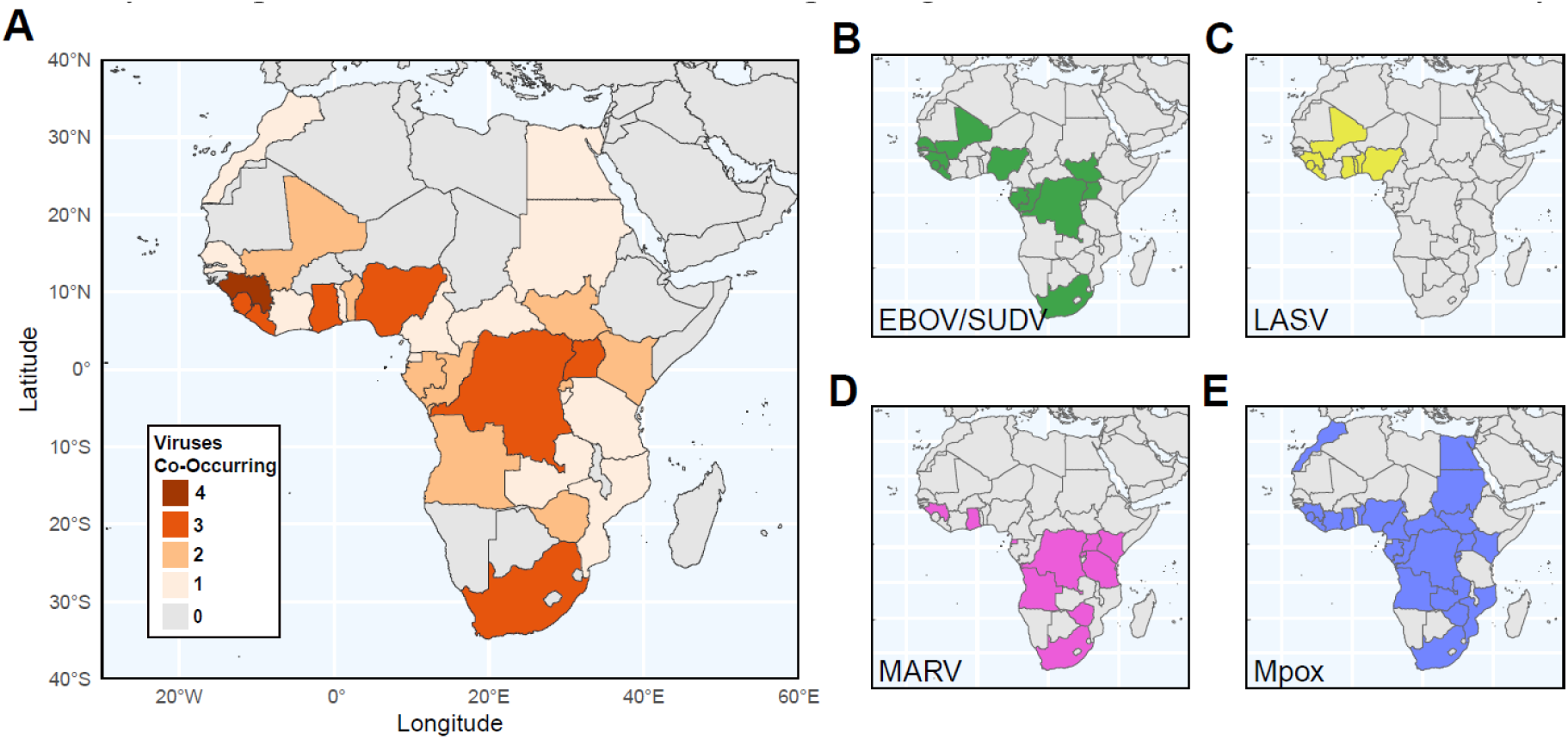
Overlapping endemic regions for VHFs and mpox in Africa highlighting the need for a single multivalent vaccine. (A) Countries where one or more of the Ebola (EBOV), and Sudan viruses (SUDV), Marburg virus (MARV), Lassa fever virus (LASV) and mpox virus have been documented to co-circulate. Geographic distribution of EBOV and SUDV (B), LASV (C), MARV (D) and mpox virus (E) mapped separately to countries with reported outbreaks or documented circulating infections.

The financial costs of outbreak responses are similarly profound. Clinical management, personal protective equipment, surveillance operations and emergency public health interventions require significant resources, while widespread illness and quarantine stalls workforce availability across essential sectors. Governments frequently rely on emergency funding and international aid, yet economic recovery remains slow where health infrastructure is already strained. Effective epidemic control relies on rapid pathogen identification, however, field diagnostics remains challenging, diagnostic capacity in many affected regions is limited, and the early clinical symptoms of VHFs overlap extensively. This complicates triage, delays implementation of appropriate control measures, such as sourcing specific vaccines and increases the risk of regional spread. A notable historical example was the 2013–2016 West African Ebola outbreak, during which initial focus on historical endemic diseases delayed recognition of the novel regional emergence of Ebola virus. This diagnostic delay contributed to widespread transmission of EBOV across multiple countries before an effective international response was mobilized (1). Similar challenges persist for other VHFs. At least 11 MARV outbreaks have been documented since 1967 (2), and other VHFs. At least 11 MARV outbreaks have been documented since 1967 (2), and more than 35 of the more diverse Orthoebolavirus species (1) have been documented. LASV, an Old World arenavirus maintained in rodent reservoirs, causes endemic disease across West Africa, with Nigeria experiencing a marked expansion of LASV affected regions from 20 to 34 of the country’s 37 states between 2018 and 2023 (2). Mpox, historically endemic in Central and West Africa, demonstrated its pandemic potential during the 2022 global outbreak with sustained transmission outside Africa. More recently, the emergence of a new lineage led to another outbreak in 2024 and another declaration of a public health event of international concern in 2024 (3). Environmental changes, including deforestation, shifting wildlife migration, climate change, and human encroachment into wildlife habitats, are further increasing spillover risk (4).

Overlapping regional outbreaks with convergent early clinical syndromes and shared operational medical challenges underscore the need for an integrated approach to disease control and prevention, particularly for frontline healthcare personnel. A single multivalent vaccine offering broad protection against multiple high-consequence zoonotic viruses would represent a transformative advance, reducing dependence on rapid diagnostics, simplifying deployment logistics, and enabling more efficient, equitable outbreak preparedness. Modified Vaccinia Virus Ankara (MVA) is an established vaccine vector with a long record of safe use in humans, including its deployment globally as a licensed mpox vaccine during the 2022 epidemic (7). With a large payload capacity for multiple different recombinant antigens (5,6), it’s natural tropism for skin and mucosal tissues, induction of humoral and cellular immunity, MVA represents an advantageous platform for vaccine development. Combined with advances in computational antigen design further enables vaccine breadth by presenting optimal structures of conserved, immunogenic epitopes capable of eliciting protective responses across diverse zoonotic viruses (7). Using the Digital Immune Optimised Synthetic Vaccine (DIOSynVax) platform, we previously developed a multivalent MVA candidate (MVA-HFVac3v1) encoding computationally designed antigens derived from SUDV and MARV glycoproteins (GPs) and the LASV nucleoprotein (NP) (6). Here, we evaluate its prophylactic, broad-spectrum protective efficacy against three distinct hemorrhagic fever viruses in simultaneous preclinical challenge studies. These “proof-of-concept” findings demonstrate that an engineered MVA-vectored vaccine can be leveraged to provide substantial cross-protection against multiple high-consequence pathogens that threaten Sub-Saharan Africa.

## Results

### Protective efficacy of DNA and MVA-HFVac3v1 vaccination regimen in lethal virus challenge studies

To evaluate the protective efficacy of the MVA-HFVac3.v1 vaccine candidate, we conducted parallel lethal challenge studies in guinea pigs using a prophylactic heterologous DNA-prime-MVA boost vaccination which was previously shown to induce durable immunity in vaccinated people. Animals received a DNA-HFVac3.v1 prime followed by two MVA-HFVac3.v1 boosts on days 28 and 56 (Figure 2A).

**Figure 2.**
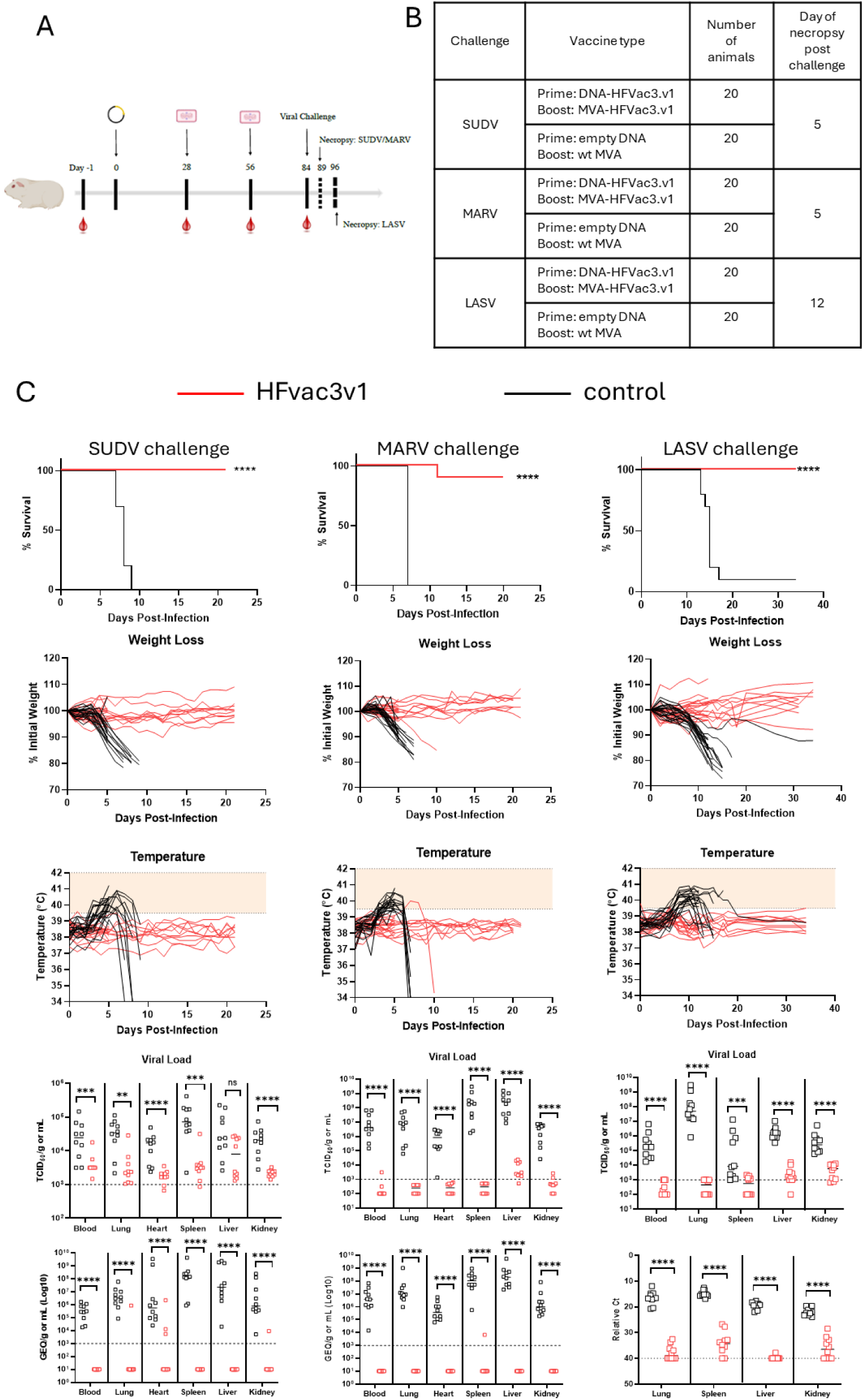
DVX-HFVac3.v1 immunisation confers robust protection against SUDV, LASV and MARV. **A**, Immunisation and timeline of challenge study. **B**, Overview of study groups. **C**, Survival outcomes in animals following virus challenge with either SUDV, MARV or LASV including overall survival, weight loss, body temperature as well as viral load in indicated tissue determined either by TCID50 or qPCR analysis. Statistical analysis of survival curves was performed with log-rank test and viral loads were analysed with non-parametric Mann-Whitney test in GraphPad prism. **** indicate p≤0.0001, *** p≤0.001, ** p≤0.01, * p≤0.05, ns p≥0.05.

The DNA and MVA-based HFVac3 boosted vaccine candidate (DVX-HFVac3.v1) was designed to encode a digitally optimized, GP sequences of SUDV and MARV respectively, together with the LASV NP, as previously described (8). Each of the three challenge experiments with SUDV, MARV or LASV included 20 vaccinated and 20 sham control animals, with equal numbers of males and females (Figure 2B). On day 84 of immunisation schedule and after the second MVA boost, animals were challenged with the corresponding heterologous virus. Half of the animals in each group were sacrificed at predetermined time points for tissue viral load analysis, while the remaining animals were monitored for survival (Figure 2B).

Across all three pathogens, the DNA primed and MVA-HFVac3.v1 boosted vaccine candidate conferred robust protection. In the filovirus challenge studies, sham controls exhibited 100% lethality, succumbing between days 7 and 9 post infection. In contrast, vaccinated animals showed 100% survival following SUDV challenge and 95% survival following MARV challenge (Figure 2C). Vaccinated animals maintained stable body weight and exhibited only minor fluctuations in temperature. Viral titers in blood, lungs, heart, spleen, liver, and kidneys were markedly reduced or below detection limit compared with sham controls. The single vaccinated animal that succumbed to MARV infection displayed weight loss and temperature changes similar to those observed in control animals.

As in humans LASV infection caused a more protracted disease course in control animals with only 5% survival, similar to human infection where LASV does not cause universally fatal disease (9). Immunisation provided complete protection, with 100% survival in the DVX-HFVac3.v1 immunized group (Figure 2C). Vaccinated animals were protected from weight loss and maintained normal ranges of body temperature. No detectable virus was found in blood, lungs, or spleen, and only near limit of detection titres were observed in liver and kidney tissues at necropsy.

### Humoral immune responses in animals prior to challenge

To identify immune correlates of protection, we evaluated humoral responses in all animals prior to lethal challenge. Blood samples were obtained prior to vaccination and 28 days after each of DNA prime and MVA boosts (therefore days -1, 28, 56 and 84 of vaccination schedule) (Figure 2A).

Due to the study size, two independent experiments were performed, one with all male and a second with all female animals. As all animals received the same vaccine batch, data from all 60 animals (30 male, 30 female) were combined for analysis.

Binding IgG responses were measured by Luminex assay, and neutralizing antibodies were assessed using a pseudovirus micro-neutralization assay (pMN). Figures 3A to C show total IgG responses against the three wild-type antigens that are phylogenetically closest to the vaccine-encoded antigens. Anti-SUDV GP and anti-MARV GP responses spanned a wide dynamic range—approximately 100-fold between lowest and highest titers. No clear correlation between antibody magnitude and survival was observed, as protected animals exhibited titers across the full range (Figure 3A–B). Notably, the single MARV-challenged vaccinee that succumbed had a median anti-MARV GP IgG response, while some survivors had lower titers (Figure 3B). Anti-LASV NP IgG responses were generally higher and less variable, approaching the upper limit of assay detection (Figure 3C).

**Figure 3.**
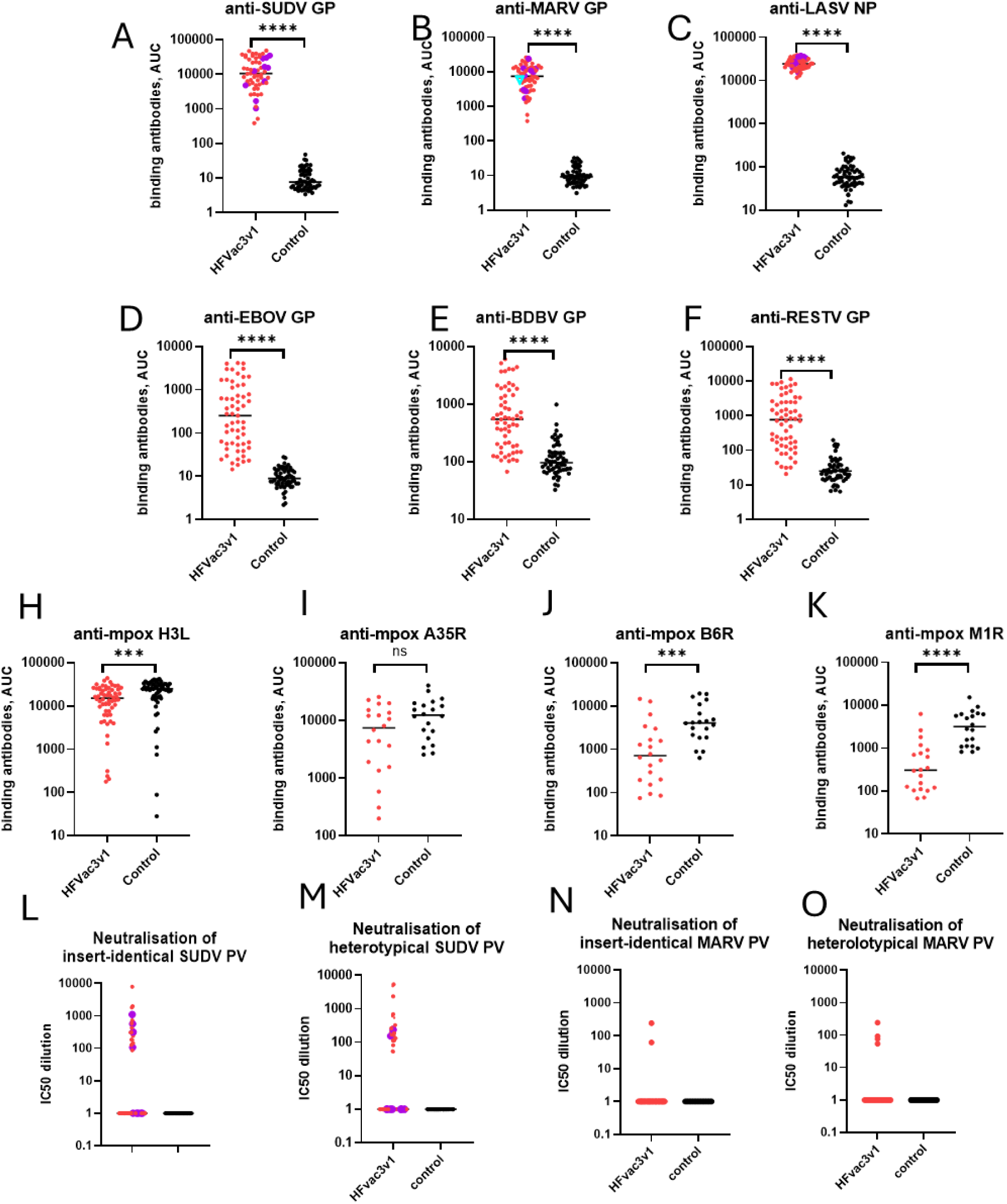
Humoral immune responses in animals following immunisation and prior to challenge with either SUDV, MARV or LASV hemorrhagic fever viruses. Panels A to K, Binding antibodies to indicated antigens measured by Luminex assay; L to O, Neutralising antibodies measured against indicated pseudoviruses by pMN assay. Vaccinated animals are shown in red, while controls are depicted in black. Every single point represents on individual animal. Notably, in panels A to C and L to M, vaccinated survivors of the respective challenges are indicated in purple, and vaccinated non-survivor in the MARV challenge is shown in cyan inverted triangle. Statistical analysis was performed with non-parametric Mann-Whitney test in GraphPad prism v10.4.2. **** indicate p≤0.0001, *** p≤0.001, ** p≤0.01, * p≤0.05, ns p≥0.05. Straight black lines indicate median value.

To assess the potential for cross-filovirus protection, we measured IgG binding to glycoproteins derived from wild-type Ebola (EBOV), Bundibugyo (BDBV) and Reston (RESTV) viruses. Most animals developed responses above background, with strongest reactivity to EBOV and BDBV GPs and lower titers to the more divergent RESTV GP (Figure 3 D-F). Luminex-based quantification of binding antibodies is a well-established and sensitive method widely used in research and clinical settings (10–13). As an orthogonal validation, we measured anti-SUDV and anti-MARV binding antibodies using a standard ELISA. ELISA results generally correlated with Luminex findings (Figure S3), although ELISA did not show a marked increase in antigen-specific titers after the second MVA boost - likely reflecting the narrower dynamic range of ELISA compared with Luminex assay. Repeated administration of the MVA-vectored vaccine at 28-day intervals did not impair boosting of either the encoded antigens or the MVA-derived H3L antigen (Figure S2D). This indicates that anti-vector immunity generated after the first MVA dose remained sufficiently low to permit productive infection of host cells and efficient antigen expression following the final booster immunization.

### Anti-mpox vector immunity

Recent studies have identified mpox antigens associated with protection against mpox disease (14,15). Anti-H3L responses reflect general anti-vector immunity, while combined responses to A35R, B6R, and M1R correlate with protection in lethal mpox challenge models. We therefore measured IgG responses to these antigens to infer the potential of DVX-HFVac3.v1 to protect against mpox. Compared with wildtype MVA, MVA-HFVac3v1 induced lower responses to H3L, B6R, and M1R, but not A35R (Figure 3H to K). Whether these differences in response either enhance or diminish protection against mpox requires evaluation in a dedicated challenge study.

### Sex-based differences in humoral responses

Because both male and female animals were included in the challenge studies, we determined if differences in antibody responses were different depending on the sex of the animals (Figure S1A–I). Female vaccinees exhibited slightly higher IgG titers against SUDV GP, MARV GP, BDBV GP, and RESTV GP (Figure S1A, B, E, F), and lower responses to mpox B6R (Figure S1I). No sex-associated differences were observed for LASV NP, EBOV GP, or mpox H3L, A35R, and M1R (Figure S1C, D, G, H, J).

### Kinetics of vaccine-induced humoral responses

To confirm the immunogenicity of the prophylactic schedule, we evaluated antibody kinetics across the three immunizations (Figure S2). Responses to all three major antigens, SUDV GP, MARV GP, and LASV NP were significantly boosted following each vaccine dose (Figure S2A to C), demonstrating strong recall responses and effective priming by the DNA-MVA regimen.

### Neutralizing antibody responses

While binding antibodies were robust in all immunised animals, the role of neutralizing antibodies (nAbs) in protection against SUDV, MARV, and LASV was less clear. Most commonly, surface-exposed glycoproteins (GP) are main targets for nAbs even though previous studies suggest that neutralizing antibodies are not required to confer protection against filoviral infection (16,17). Studies of LASV survivors in endemic regions have indicated that intravirion protein LASV NP and not surface-exposed GP is the key protective antigen (18,19). Interestingly, in DVX-HFVac3.v1, LASV NP alone was sufficient to confer 100% protection against lethal LASV challenge.

For SUDV and MARV, we quantified neutralizing responses prior to challenge using pseudovirus (PV) assays incorporating either insert-identical or naturally occurring wildtype GPs. Neutralizing anti-SUDV responses spanned a 100-fold range of IC50 values and were detectable in approximately half of all vaccinated animals (43% for insert-identical and 47% for heterotypic SUDV GPs) (Figure 3L to M). Among vaccinated survivors of SUDV challenge, fewer than half had detectable nAbs, indicating that nAbs were not required for protection. Animal sex did not influence the frequency or magnitude of SUDV-specific nAb responses (Figure S2K to L). Analysis of nAb kinetics showed that among animals that developed detectable nAbs, approximately half did so after the first MVA boost, with the remainder responding after the second boost. This pattern was independent of sex (Figure S2E). As for neutralizing responses to MARV, nAbs against both insert-identical and heterotypic GPs were undetectable in most animals (Figure 3N to O).

### Cellular immune responses in animals prior to lethal virus challenges

The selection of the LASV NP as our lead vaccine antigen was based on our field studies in Lassa fever endemic communities in Africa (19), where NP-specific T cell responses were identified as key correlates of protection. Given the high cross-reactivity of NP-specific T cells across LASV lineages (18), we evaluated cellular immunity in vaccinated animals by stimulating PBMCs with pools of 15mer overlapping peptides spanning the insert-matched LASV NP sequence. T cell responses to SUDV and MARV GP antigens have also been reported previously (20,21). Therefore, PBMCs were independently stimulated with overlapping peptide pools corresponding to insert-matched EBOV GP or MARV GP. PBMCs collected two weeks after the first and second MVA boosts (days 42 and 70) were assessed using an in-house optimized IFNγ ELISpot assay (22).

LASV NP elicited strong peptide-specific IFNγ responses in 90% of animals on day 42 (27 out of 30 tested samples) and 76% on day 70 (22 out of 29 tested samples), with no substantial increase in magnitude or frequency after the second MVA boost (Figure 4). MARV GP peptides also induced robust T cell responses, with 74% of animals responding on day 42 (20 out of 27 tested samples) and 82% on day 70 (23 out of 28 samples). EBOV GP-specific responses were detectable in a smaller proportion of animals with 44% on day 42 (11 out of 25) and 27% on day 70 (7 out of 26), consistent with previous reports (20). T cell response magnitude and frequency were not influenced by sex (Figure S4).

**Figure 4.**
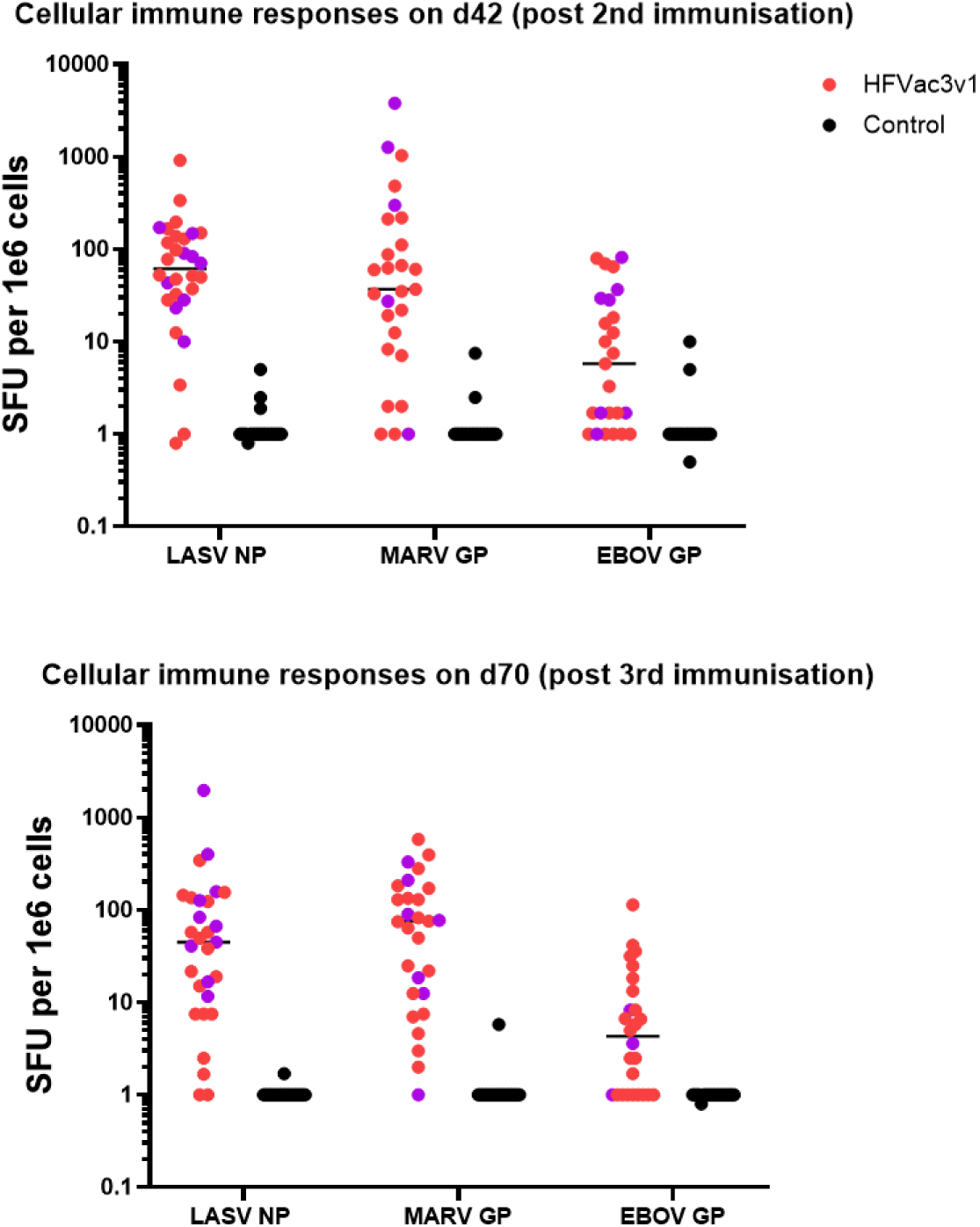
Cellular immune responses in animals prior to challenges with either SUDV, MARV or LASV. PBMCs were stimulated with indicated insert-matching peptide pools for LASV NP, MARV GP or EBOV GP either post second (on day 42) or post third immunisations (on day 70). Results are represented as spot forming units (SFU) per 1 million cells. Minimum of 10 samples per group per animal sex were analysed. Purple dots indicate responses in confirmed survivors of respective challenges whereas red combine both confirmed survivors in other challenges and animals culled at necropsy time point before survival could be ascertained.

## Discussion

The robust simultaneous protection achieved against SUDV, MARV and LASV by the DVX-HFVac3.v1 vaccine candidate demonstrates the potency of the computationally designed antigens delivered within a single MVA-vectored platform. Despite disease developing in one animal in the MARV challenged group, the overall findings provide further insight into immune correlates relevant for next generation vaccines targeting filoviruses and Old World arenaviruses.

Compared with existing rVSV and adenoviral vectored approaches, an MVA backbone offers practical advantages for population safety, manufacturing/logistics, and economic efficiency in African settings. Importantly, MVA has an established human safety profile, including use during the 2022 mpox response, with favourable tolerability across diverse groups (immunocompromised individuals, children, pregnant people) and minimal risk of vector replication or neurotropism, considerations that may constrain rVSV and some adenoviral platforms in mass campaigns. Secondly the payload capacity of MVA enables multivalent expression of computationally designed, conserved antigens from SUDV, MARV, and LASV within a single vaccine vector closely related to a licensed product, simplifying chemistry, manufacturing, and controls (CMC), reducing tech-transfer complexity, and leveraging existing fill-finish and cold-chain pathways already used for mpox vaccines (23,24). Third, a combined VHF vaccine has the potential be deployed prophylactically to high-risk individuals such as health care workers or populations with a high likelihood of seasonal recurrence. Additionally, an accelerated immunisation protocol (not examined here) may be deployed early upon presentation of VHF-like illness when clinical overlap and limited diagnostics delay pathogen identification, thereby reducing time-to-immunization, and stockpile fragmentation. The operational burden of maintaining multiple pathogen-specific products is significant and economically, consolidating procurement and distribution into one or fewer products lowers per-dose costs through scale efficiencies, decreases wastage from expiring, pathogen-specific inventories, and offers advantages in streamlining immunisation policies, practice and mitigating consequences of exposure risk for key frontline health care staff. Collectively, these attributes argue that single, computationally designed multivalent vaccine can deliver broader public-health benefit per logistical unit than other single-disease vaccine comparators, especially where diagnostics are constrained, and rapid, equitable coverage is essential.

Human phase 3 efficacy trials for targeted pathogens are notoriously difficult due to the sporadic and unpredictable nature of outbreaks, logistical challenges in remote regions, as well as the difficulties in long-term volunteer retention in these regions. Consequently, regulatory agencies have relied heavily on animal rule pathways, granting conditional marketing authorization to rVSV-ZEBOV (Zerbevo) and “under exceptional circumstances” approval to the Zabdeno/MVA-BN-Filo regimen both licensed solely for Ebola Zaire and based initially on animal efficacy data (25,26). Among available preclinical models, outbred guinea pigs offer advantages over mouse models due to their more human-like disease pathogenesis and immune responses, while nonhuman primate studies, though immunologically closest to humans, are constrained by cost, small sample sizes and limited representation of sex and age (27). Previous EBOV vaccine development demonstrated strong correlation between total GP-specific IgG and survival in guinea pigs, mirroring findings in nonhuman primates (16).

Specific threshold immune correlates of protection were not possible to discern in this study because of the high level of vaccine induced protection (in all but one animal). In the SUDV challenge model, all vaccinated animals developed binding antibodies to SUDV GP, though neutralizing antibody (nAb) titers varied widely. Protection was achieved even in animals lacking detectable nAbs, indicating that neutralization is not essential for SUDV protection. This aligns with earlier studies of Ad5-based EBOV vaccines in guinea pigs and rVSV-EBOV in nonhuman primates, where survivors often exhibited highly variable or undetectable nAb levels (16,17). The DVX-HFVac3.v1 vaccine regimen induced only modest SUDV GP-specific T cell responses, suggesting that humoral immunity, specifically total binding IgG is contributing to protection against SUDV. Similarly, DVX-HFVac3.v1 vaccinated animals developed a broad range of MARV GP-specific IgG titers, yet survival did not correlate with antibody magnitude. The single non survivor exhibited median IgG levels, while some protected animals had titers an order of magnitude lower. Neutralizing antibodies were largely undetectable against both insert-identical and heterotypic MARV pseudoviruses, indicating that neutralization is not a correlate of protection as shown in several studies (28,29). Notably, MARV GP elicited strong T cell responses in most vaccinated animals, suggesting that cellular immunity may contribute to protection in the guinea pig model.

Immune correlates of protection against LASV were similarly found to correlate with NP responses in LASV NP immunised animals. The choice of LASV NP was guided by field studies in endemic regions, where LASV NP-specific T cell responses correlate with survival (18,19) as well as previous challenge studies (30). Consistent with this, DVX-HFVac3 vaccinated animals exhibited strong LASV NP-specific T cell responses and achieved 100% protection against lethal challenge with heterologous LASV. Although LASV NP does not induce neutralizing antibodies, it generated robust binding IgG responses, suggesting that both cellular and humoral immunity may contribute to viral control. These findings reinforce the suitability of LASV NP as a lead LASV vaccine antigen.

A single multivalent vaccine capable of protecting against unpredictable outbreaks of high consequence pathogens would provide substantial public health and economic benefits for Sub-Saharan Africa. Ebola, Marburg, and Lassa fever are consistently listed by WHO as priority diseases due to their epidemic potential and limited countermeasures (31). The WHO has also emphasized the need for multivalent filovirus vaccines (32) given the overlapping clinical presentations and cocirculation of these pathogens. The DVX-HFVac3 vaccine provides a compelling proof-of-concept that a single vaccine can protect against at least three regionally overlapping diseases, SUDV, MARV and LASV (Figure 1), with clinically indistinguishable symptoms. Our serological data combined with findings by others demonstrating MVA-mediated protection against mpox infection suggest that DVX-HFVac3 may protect against 4 different viruses endemic in Sub-Saharan Africa (33).

Economic gains are several fold. Firstly, a multivalent “4-in-1” vaccine would reduce manufacturing and distribution costs and improve vaccine uptake by healthcare workers and communities in endemic regions. Such a vaccine would be particularly advantageous in remote regions where rapid diagnostic differentiation between hemorrhagic fevers is challenging. For healthcare workers and first responders, a single prophylactic vaccine could provide broad protection during early outbreak stages when pathogen identity is uncertain. Beyond health benefits, the economic implications are substantial. High consequence viral outbreaks impose severe societal costs through quarantines, travel restrictions, and workforce losses. Modelling of Lassa fever alone suggests that even a modestly effective vaccine (70% efficacy, 32.5% coverage) administered over three years could avert approximately 80 million international dollars in societal costs and nearly 800 million dollars in value of statistical life losses over a decade (34). A multivalent vaccine would therefore be highly attractive to governments, global health organizations, and investors.

Importantly, future studies are required to identify optimal immunization regimens in different setting such as prophylactic immunization of front-line health care workers populations living in regions frequently impacted by hemorrhagic fever outbreaks, versus the current practice of outbreak response immunisation, in which single, rapid immunisation formulations are highly desired. This study used an immunisation schedule to address prophylactic immunisation of front-line health care workers. Future studies plan to explor optimal single dose or rapid prime-boost schedules for outbreak response protocols. Thus, to enhance durability of immunity in high risk front line healthcare workers, we employed a heterologous DNA prime MVA boost regimen, which has previously been shown to generate longer lasting responses than MVA only regimens such as JYNNEOS (35). Clinical studies are planned to determine the optimal regimen for durability of immune responses, scalability, and affordability.

## METHODS

### Production of vaccines

Design of DNA-HFVac3v1 and MVA-HFVac3v1 is described elsewhere (8). DNA-HFVac3.v1 and empty pDNA was purified using the EndoFree Plasmid Mega kit (Qiagen, Hilden, Germany). MVA-HFVac3v1 and wild-type MVA were propagated in suspension AGE1.CR.pIX cells (ProBioGen).

### Ethics statement

All animal work was conducted in compliance with the guidelines established by the Canadian Council on Animal Care, as approved by the Animal Care Committee at the Canadian Science Center for Human and Animal Health (CSCHAH: animal use document H-22-006 and H-22-009). Guinea pigs were monitored by registered Animal Health Technicians at the National Microbiology Laboratory (NML) (Winnipeg, MB, Canada) throughout the experiment and were provided food and water ad libitum. Animals were group housed under controlled laboratory conditions, including a 12-hour light/dark cycle, a temperature range of 21–22°C, and humidity levels of 30–40%. Animals were acclimatized for 7 days before the beginning of all experiments.

### Viruses and Cells

Guinea pig adapted (GPA) SUDV, MARV, and LASV strains were previously developed by researchers from the National Microbiology Laboratory (NML) (36,37). Vero cells were maintained in MEM (Gibco, 10370-021) supplemented with 5% heat inactivated Bovine Growth Serum (BGS, Cytivia Hyclone, SH30541.03), 2mM L-Glutamine (Gibco, 25030-081), and penicillin and streptomycin (P/S; 100 Units/mL and 100 µg/mL respectively; Gibco, 15140-122). VeroE6 cells were maintained in DMEM (Gibco, 11965-092) with the same supplementation.

### Vaccine regimens

Adult outbred male and female Hartley guinea pigs (Cavia porcellus) were purchased from Charles River Laboratories (Wilmington, MA, USA) and housed in a biosafety level 2 (BSL2) animal facility during the course of immunization at the NML. Animals were subcutaneously implanted with a BMDS IPTT-300 programmable temperature transponder (BMDS, Seaford, DE, USA) into the intra-scapular region. In each viral challenge group (SUDV, MARV, or LASV), 40 guinea pigs were divided into groups of 20 (10 males/10 females) and were randomly assigned into each of the control and vaccination arms for a total of 120 animals. All experimental manipulations were conducted on sedated animals (3–5% inhalational isoflurane (Baxter, CA2L9108) maintained in medical oxygen). Prime vaccination was completed by intradermal (ID) delivery of 100 ug of DNA in 100 uL PBS (Corning, 21-040-CV) using the PharmaJet TropisID Injector (Golden, Colorado, USA) with each injection delivering 100 uL per hind leg of DNA HFVac3.v1vaccine or control DNA empty for the vaccinated and control groups respectively. Subsequent boosts at 28 and 56 dpv consisted of 2 x 107 PFU of MVA HFVac3.v1 or MVA WT empty for the vaccinated and controls groups, respectively, by intramuscular injection. Serum was collected at each vaccination timepoint (0, 28, and 56 days post vaccination (dpv)), and on the day of viral challenge (84 dpv, 0 days post infection (dpi)).

### Luminex binding antibody assay

Antigens were covalently coupled to Luminex MagPlex® beads using EDC/S-NHS chemistry in MES buffer (pH 6.0), following a modified protocol from the xMAP® Cookbook (5th Edition, Luminex). Coupling was performed with 12.5 µg of recombinant protein per 1.25×10⁷ beads. The following recombinant proteins were used: Sudan virus glycoprotein (referred to as SUDV GP, IBT Bioservices, #0502-001), Ebola virus Zaire glycoprotein (ZEBOV GP; IBT Bioservices, #0501-001), Musoke Marburgvirus glycoprotein (MARV GP, IBT Bioservices, #0503-001), Ravn virus glycoprotein (RAVV ΔMuc GP, IBT Bioservices, #0513-015), LASV nucleoprotein lineage III full length (LASV NP; Zalgen, #LASV-R-0031), MPXV A35R protein Stratech, #40886-V08H-SIB,), MPXV B6R protein (Stratech, #40902-V08H-SIB), MPXV H3L protein (Sino Biologicals, #40893-V08H1), MPXV M1R protein (Stratech, #40904-V07H-SIB). Coupled beads were stored in PBS with 1% BSA and 0.05% sodium azide at 4°C.

Serum samples were diluted (1:100, 2-fold, 8-point series) in PBS with 1% milk powder and incubated with coupled bead mixes in black 96-well plates for 1 h at 37°C, 600 rpm. After washing in PBS-T (PBS + 0.05% Tween-20), beads were incubated with PE-conjugated goat anti-mouse IgG (H+L) cross-adsorbed (Invitrogen, #P852) or donkey anti guinea pig F(ab’)2 fragment (Jackson ImmunoResearch, #706-116-148) in Luminex assay buffer (PBS with 1% BSA, 0.05% NaN₃, 0.5% PVA, 0.8% PVP) for 30 min at 37°C. Plates were washed, resuspended in PBS-T, and read on a Luminex-200 analyzer with the default setting of measuring the fluorescence intensity of 50 beads per colour/target. Results are given as mean fluorescence intensity (MFI). The MFI on each plate was normalised to a positive reference control serum. MFI was plotted against the log10(dilution factor) and the AUC was calculated for each binding curve. As NIBSC standards were not included as they are not commercially available for murine or guinea pig sera. Sensitivity and selectivity of the Luminex assay for these antigens is described elsewhere (8).

### Lethal Challenge with guinea pig-adapted virus strains and Sample Collection

Twenty-eight days after the third immunization according to the immunization schedule in Figure 2, animals were transferred to the BSL4 facility at the NML and challenged with 1000x median lethal dose (LD50) of either GPA-SUDV Boniface, 1000x LD50 of GPA-MARV Angola, or 100x LD50 of GPA-LASV Josiah via the intraperitoneal route (IP). Daily animal checks were conducted which included recording temperature, weight, and clinical score for all animals up to 20 dpi for GPA-SUDV and GPA-MARV animals, and 34 dpi for GPA-LASV animals. Oral and rectal swabs were sampled for all animals at 3 and 5 dpi (GPA-SUDV and GPA-MARV groups). For GPA-SUDV and GPA-MARV challenge groups, half the animals in each group were euthanized at 5 dpi and necropsied for collection of blood, serum, lung, heart, spleen, liver, and kidney. For the GPA-LASV challenge groups, half of the animals in each group were euthanized at 12 dpi and necropsied for collection of blood, serum, lung, liver, spleen and kidney. The procedure for euthanasia at 5 dpi or 12 dpi, as well as at later dates for animals meeting humane endpoint criteria was exsanguination via cardiac puncture while under deep anaesthesia followed by a thoracotomy.

### Live Virus Assay

Infectious viral titres were measured in blood and tissue samples (lung, heart, spleen, liver, and kidney) which were collected during necropsy. Tissue samples were placed in cryovials, weighed, and flash frozen at -80°C until further analysis. Tissues were homogenized in 1 mL viral growth media (MEM supplemented with 1% BGS and P/S (100 Units/mL and 100 µg/mL respectively; Gibco)), along with a 5 mm sterile stainless-steel bead using a Bead Ruptor Elite Tissue Homogenizer (Omni) set to 4 m/s for 30 s. Homogenates were clarified by centrifugation at 1500 x g for 10 minutes, and ten-fold dilutions of tissue homogenates and whole blood were made in viral growth media. Dilutions (100uL of each) were added to triplicate wells of Vero (GPA-SUDV and GPA-LASV) or VeroE6 (GPA-MARV) in 96-well plates. The cells were incubated at 37°C with 5 % CO2, and cytopathic effect was read at day 14 (GPA-SUDV), day 10 (GPA-MARV) or day 7 (GPA-LASV). TCID50 values per gram of tissue were calculated using the Reed and Muench method (38).

### RNA Load Analysis

The tissues that were collected from GPA-MARV and GPA-SUDV infected animals at 5 dpi or from GPA-LASV-infected animals at 12 dpi were placed in cryovials with 500 uL of RNAprotect Tissue Reagent (Qiagen, 1018087). After overnight incubation at 4°C, RNAprotect was removed and the tissues were weighed. Tissues were then homogenized in 600 uL Buffer RLT Plus (Qiagen, 1048449) with a 5 mm stainless-steel beads using a Bead Ruptor Elite Tissue Homogenizer (Omni). RNA was extracted from the homogenized tissues using the RNeasy Plus Mini Kit (Qiagen, 74136) and from whole blood using Qiagen QIAamp Viral RNA Mini Plus kit (Qiagen, 52906) followed by MagMax Nucleic Acid Isolation Kit on the KingFisher Apex prime platform (ThermoFisher Scientific). Viral genomes of GPA-SUDV and GPA-MARV were detected with a QuantStudio 5 Real-Time PCR System instrument (Applied Biosystems) using TaqMan 1-step Fast Virus Mix (Applied Biosystems, 4444432), with the RNA polymerase gene as the target. Cycling parameters were 25°C for 2 min, 53°C for 10 min, 95°C for 2 min, followed by 40 cycles of 95°C for 3 s and 60°C for 30 s. Primer and probe sequences used in the reactions were as follows: SUDV-forward: 5’-CAGAAGACAATGCAGCCAGA-3’; SUDV-reverse: 5’-TTGAGGAATATCCCACAGGC-3’; SUDV-probe: 6-carboxyfluorescein (FAM)-CTGCTAGCTTGGCCAAAGTCACAAG-black hole quencher (BHQ); MARV-forward: 5’-GCAAAAGCATTCCCTAGTAACATGA-3’; MARV-reverse: 5’-CACCCCTCACTATRGCGTTYTC-3′; MARV-probe: 6-carboxyfluorescein (FAM)-TGGCACCAYAATTCAGCAAGCATAGG-black hole quencher (BHQ). Qualitative molecular detection of LASV was performed using an in-house, validated diagnostic RT-PCR test with primers and probe targeting the glycoprotein gene: LASV-forward: 5’-ATGGCTTGTTTGAAGTCRAA and LASV-reverse: 5’-TGACCAGGTGGATGCTAATTGA, and LASV-probe 5’-(FAM)-CATGTCACAAAATTCTTCATCGTGCTTCTCA. Reactions were carried out on a QuantStudio 3 Real-Time PCR System instrument (Applied Biosystems) using the QuantiTect Probe RT-PCR kit (Qiagen) according to the manufacturer’s specifications.

### Detection of anti-SUDV GP and anti-MARV GP specific antibodies by ELISA

A 96-well enzyme-linked immunosorbent assay (ELISA) low binding plate (Nunc F-96 Well Plate, Non-Treated Surface, Thermo Fisher Scientific cat #269620) was coated overnight at 4°C with 200 ng per well of baculovirus-expressed SUDV glycoprotein (GP) made in house using published methods (39) or MARV GP (IBT bioservice cat # 0506-015) in PBS. The plates were washed 5 times in wash buffer (PBS, 0.1% Tween 20) and blocking buffer was added to each well and incubated at 37°C for 1 hour (5% skim milk powder, Difco cat#232100 in PBS). Guinea pig sera collected at 0, 28,56 and 84 dpv were diluted 1:100 in blocking buffer followed by two-fold dilutions a total of 8 times. 50µL of each sample was added to the ELISA plates and incubated for one hour at 37°C. Plates were washed as before and 50µL Goat anti-guinea pig IgG –HRP secondary antibody (1:1000, KPL cat# 14-17-06) was added for one hour at 37°C. The plates were washed and 50µl of TMB (Life Technologies cat#002023) was added to each well. After 15 minutes of incubation in the dark at room temperature, 50 µl of 1M sulfuric acid (H2SO4) was added, and the plates were analyzed on a Synergy HTX (BioTek) microplate reader at 450 nm wavelength.

### Production of pseudotyped lentiviral vectors

Lentiviral pseudotypes were produced by transient transfections of HEK293T/17 cells (4x10^5^ cells/6-well) with lentiviral packaging plasmids p8.91-gag-pol and pCSFLW-firefly luciferase (64,65) and the DNA plasmid encoding the respective glycoprotein (SUDV-GP Gulu 2000, GenBank: NC_006432; SUDV-GP Uganda 2022, GenBank: OQ672950.1; ZEBOV-GP Yambuku 1976, GenBank: NC_002549; MARV-GP Musoke 1980, GenBank: NC_001608; proprietary EBOV T2-4 GP gene; proprietary MARV T2-11 GP gene) using the FuGENE®-HD Transfection Reagent (Promega).

Supernatant was harvested after 48 h, filtered through a 0.45 µm filter, aliquoted, and immediately stored at -80°C. Each aliquot was defrosted only once and discarded after use. Pseudotyped viruses were titrated on HEK293T/17 cells.

### Pseudotype microneutralisation assay

Pseudotype-based microneutralization assays (pMN) were performed as described previously (66). Briefly, serial dilutions (1:2) of serum (1:33 start) were incubated with SUDV-GP Gulu 2000, ZEBOV-GP Yambuku 1976, MARV-GP Musoke 1980, insert-identical T2-4 EBOV GP or insert0identical MARV T2-11 GP bearing lentiviral pseudotypes, respectively for 1 h at 37 °C and 5% CO_2_ in white 96-well culture plates. HEK293T/17 cells (1.5 × 10^4^) were then added per well and plates incubated for 48 h at 37 °C and 5% CO_2_ in a humidified incubator. Bright-Glo (Promega) was then added to each well and luminescence read after 5-min incubation using the GloMax microplate reader (Promega). Experimental data points were normalised to 100% and 0% neutralization. Normalised responses were plotted against the dilution factor and fitted to the nonlinear model ‘log(inhibitor) vs normalised response – Variable slope’.

### ELISpot in guinea pig PBMCs

During the vaccination schedule, 3 mL of whole blood was collected at 42 and 70 dpv into K2 EDTA (K2E) vacutainers (BD, 467856) and processed for collection of peripheral blood mononuclear cells (PBMCs). The blood was mixed 1:1 with Hank’s Balanced Salt Solution (HBSS; Gibco, 14175) and added to a SepMateTM-15 tube (StemCell Technologies, 85415) containing 4.5 mL of Ficoll-PaqueTM Plus (Cytivia, 17144002). The mixture was centrifuged at 1200 rcf for 10 minutes, and the top layer was decanted into a clean conical tube which was topped up to 25 mL with R10 medium (RPMI 1640 media (Cytivia, SH30027.1) supplemented with 10% heat inactivated fetal bovine serum (FBS; Corning, 35-015-CV) and 1% P/S). Cells were washed by pelleting at 450 rcf for 5 minutes, followed by resuspension in R10 media and pelleting again. The cells were then resuspended in 1 mL R10 media and passed through a 70 µm cell strainer (Fisherbrand, 22-363-548) into a new tube. Cell counting was accomplished by mixing 20 µL of cells 1:1 in Acridine orange (5 µg/mL; Invitrogen, A3568) and Propidium iodide (100 ug/mL; Invitrogen, P3566) in PBS and examining on a Cellometer (Nexcelom). Afterwards, cells were pelleted at 450 rcf for 5 minutes, and resuspended in freezing media (cold FBS with 10% DMSO (Sigma, D2650)). Cells were then frozen in 1.2 mL Cyogenic vials (Corning, 430487) at -80°C overnight in a Corning CoolCell chamber, and then stored in liquid nitrogen dewars.

ELISpot assay was executed as published earlier with modifications (22). PBMCs were thawed, counted and incubated at the same cell density with overlapping 15-mer peptide pools of either LASV NP, EBOV GP or MARV GP antigens for 20 hours at 37 °C and 5% CO_2_ in a humidified incubator. Capture and detection antibodies were produced by GenScript and titrated to establish optimal concentrations for the assay performance. The assay was developed with ELISpot conjugate: Streptavidin-ALP and BCIP/NBT-plus substrate for ALP (Mabtech). The plates were dried, scanned and analyzed on ELISpot reader S6 Ultra M2 (ImmunoSpot).

### Statistical analysis

Data were analyzed using GraphPad Prism software version 10.4.2. Non-parametric tests were used to investigate statistical differences. In general, differences between two groups were tested for statistical significance using the Mann-Whitney test. For the analysis of the kinetics of antibody induction, the Wilcoxon paired signed rank test was used.

---

**Figure S1.**
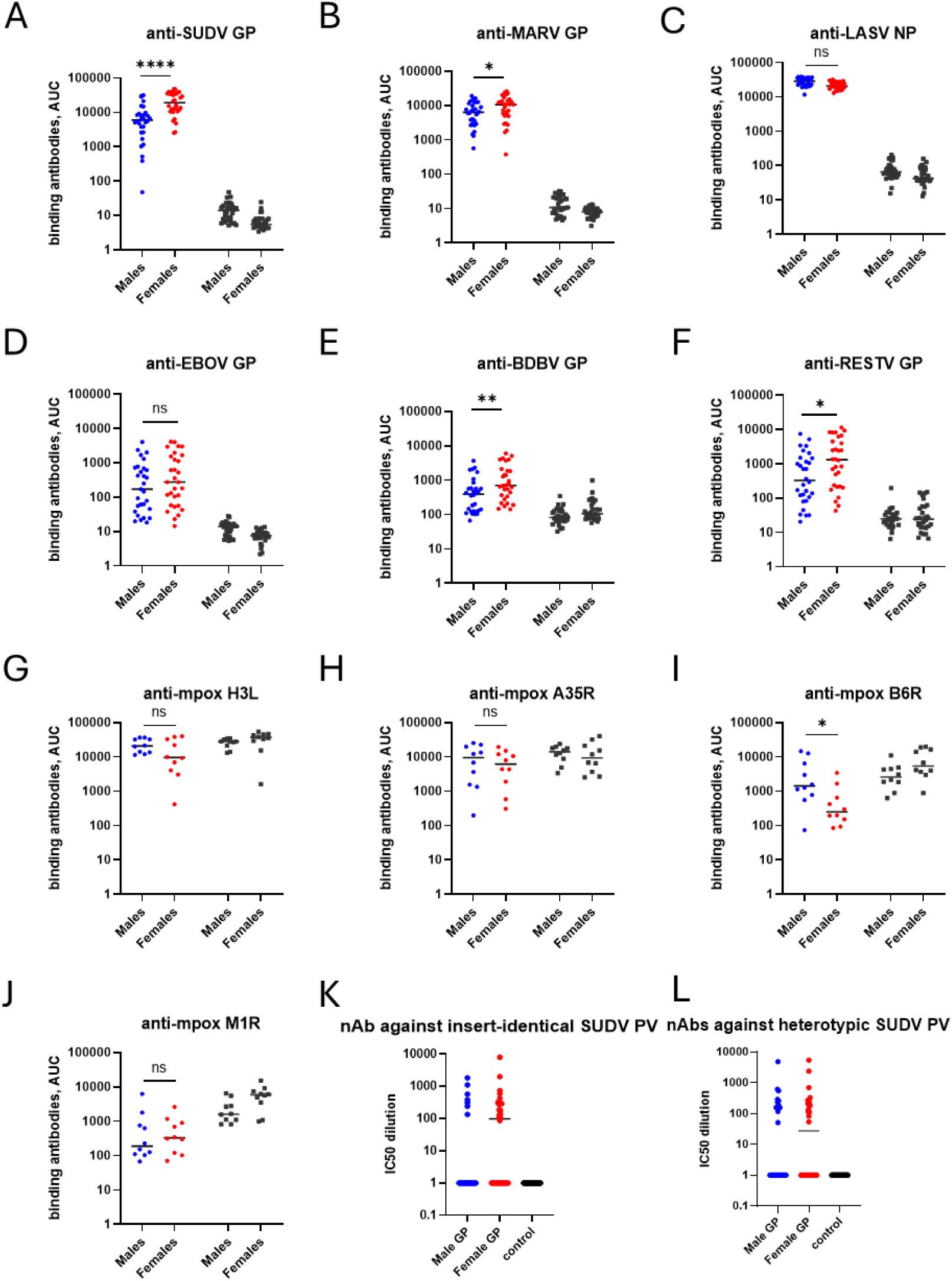
Humoral immune responses against indicated antigens following immunization with DVX-HFVac3.v1 vaccine and prior to challenge with either SUDV, MARV or LASV hemorrhagic fever viruses in male vs female animals. Panels **A-J**, Binding antibodies to indicated antigens measured by Luminex assay; **K-L**, Neutralising antibodies measured against indicated pseudoviruses by pMN assay. Statistical analysis was performed with non-parametric Mann-Whitney test in GraphPad prism v10.4.2.

**Figure S2.**
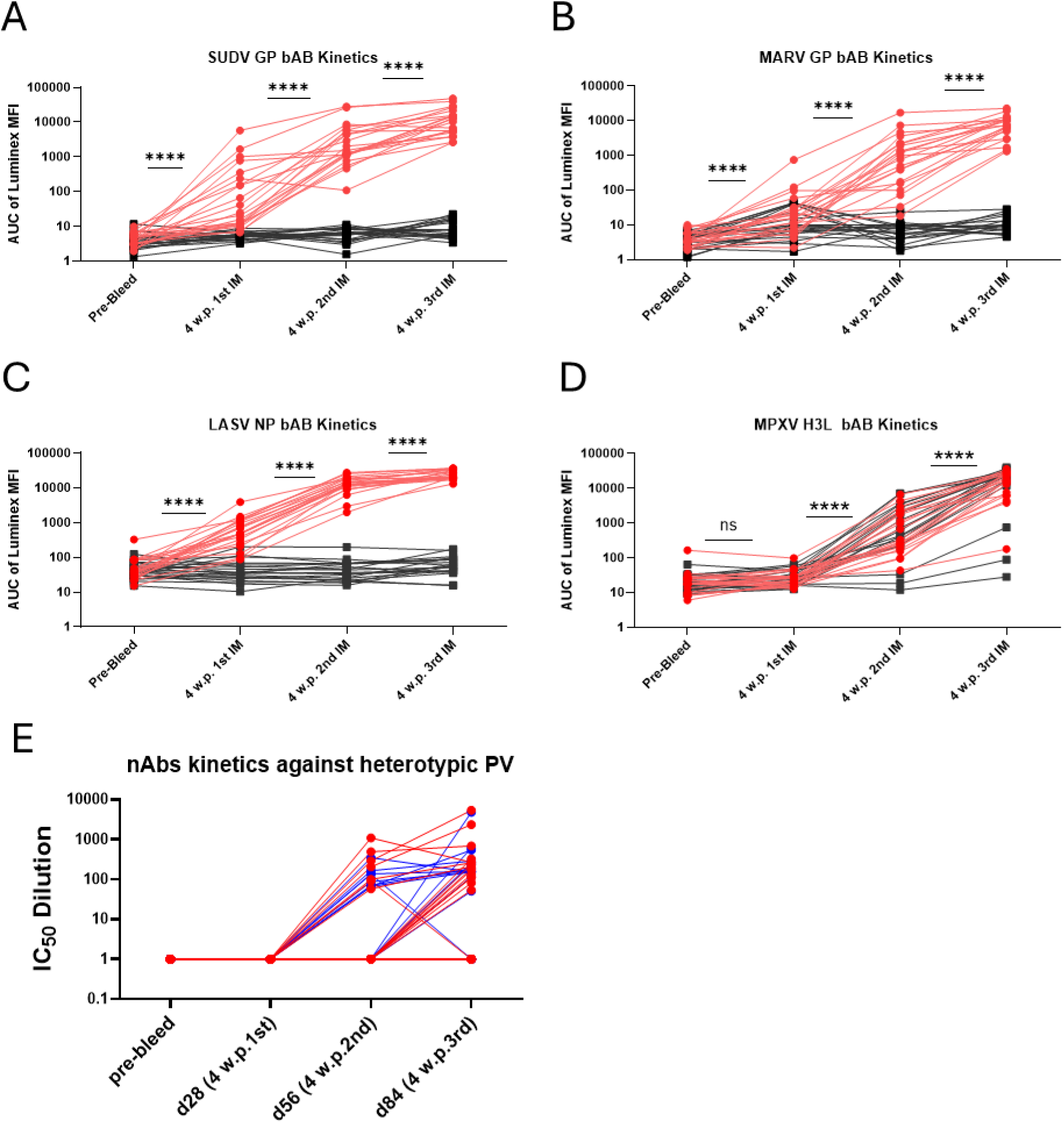
Kinetics of humoral responses against indicated antigens as measured for binding antibodies by Luminex (panels **A-D**) or for neutralising antibodies by pMN methodologies (**E**). For neutralising antibodies, comparison is performed between male (in blue) and female animals (in red). Statistical analysis for panels A-D was done in GraphPad Prism 10.4.2 by using Wilcoxon matched-pairs signed rank test.

**Figure S3.**
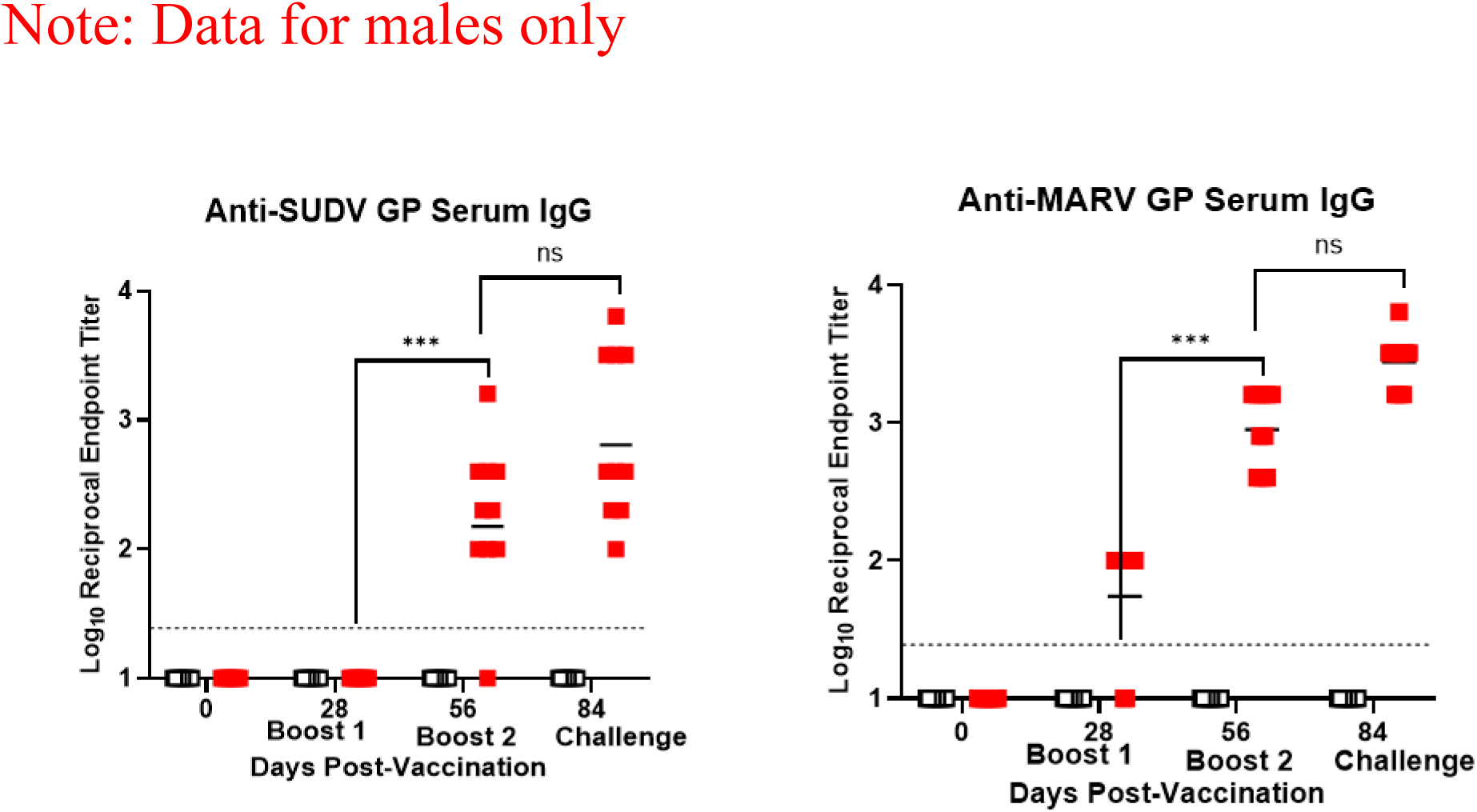
Anti-SUDV and MARV GP ELISA in animal serum of male guinea pigs post vaccination with DVX-HFVac3.v1 vaccine candidate. Serum samples were taken at the indicated time points. Statistical analysis was done in GraphPad Prism 10.4.2 using Mann-Whitney test.

**Figure S4.**
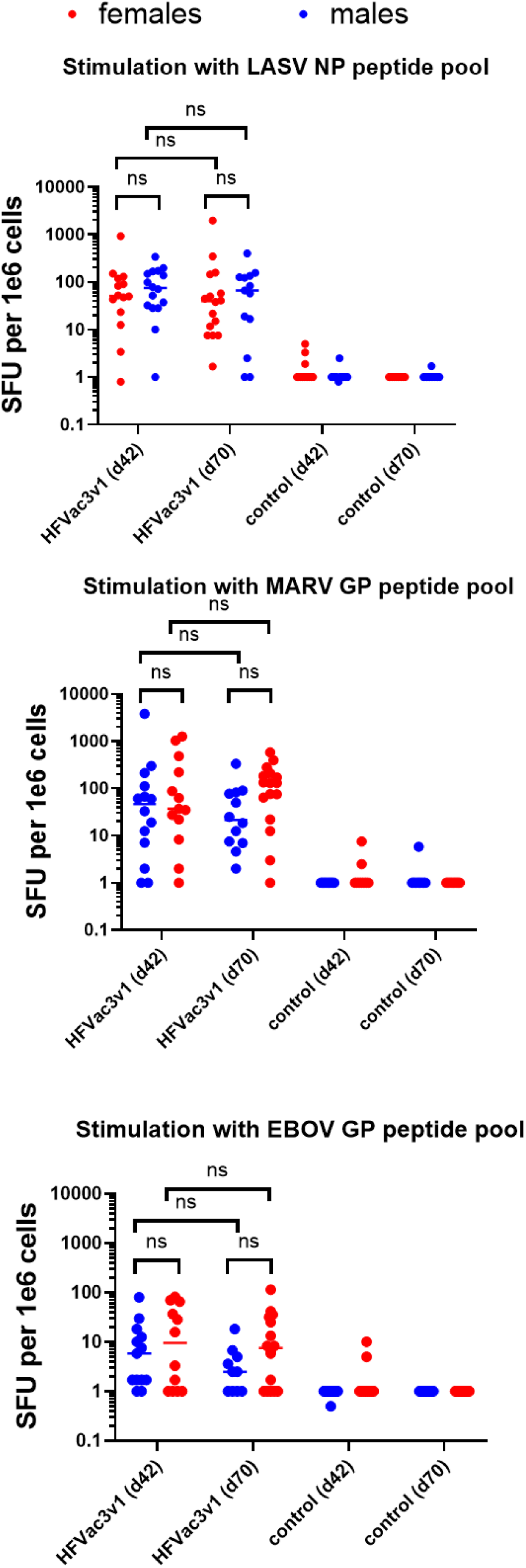
Cellular immune responses stimulated with indicated peptide pools and at the indicated time points (day 42 and day 70) in male vs female DVX-HFVac3.v1 vaccine recipients.

