## Supplementary figures and images for "Simultaneous broad protection against Ebola Sudan, Marburg and Lassa viruses conferred by a DNA primed MVA-vectored multivalent vaccine"

### Figure S1

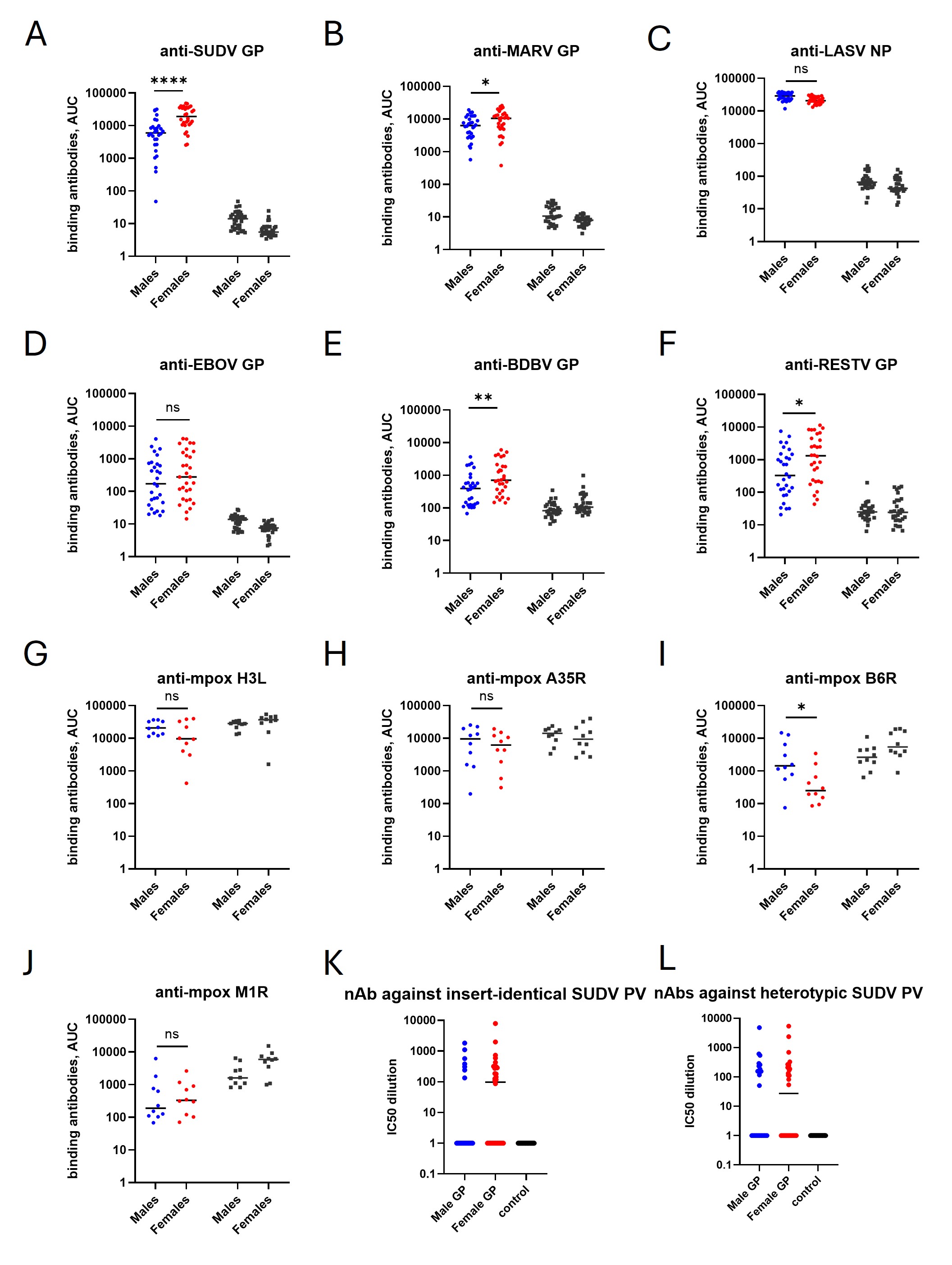

### Figure S2

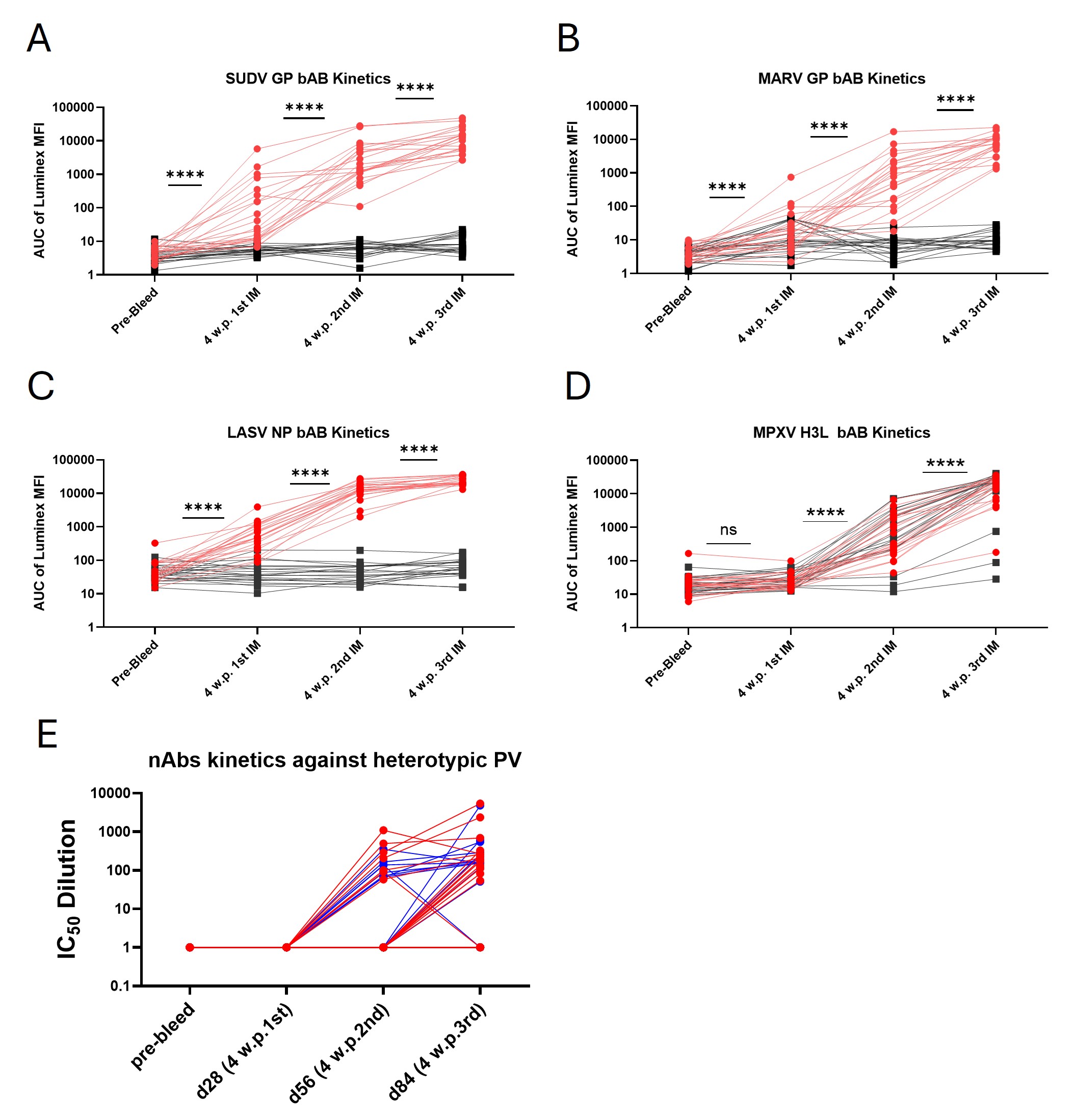

### Figure S3

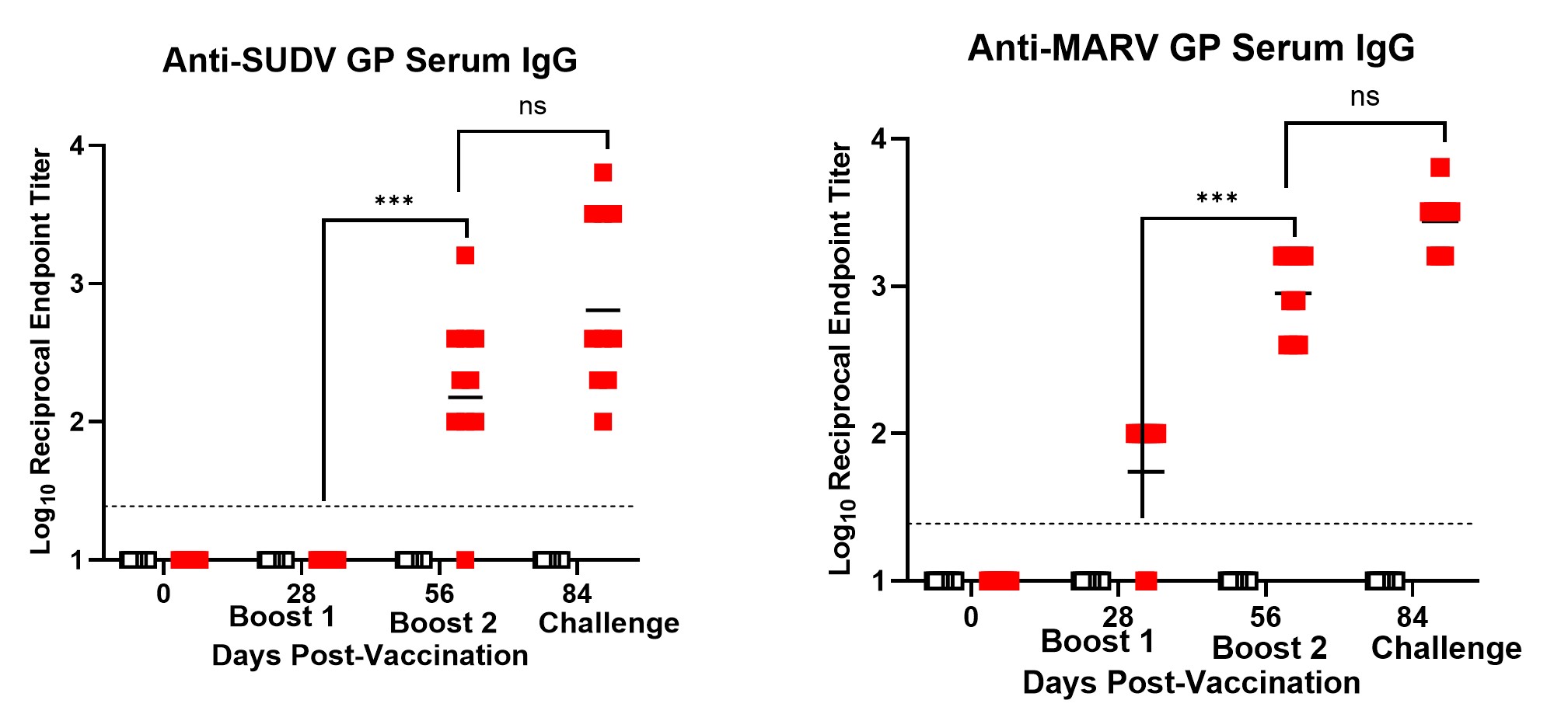

### Figure S4

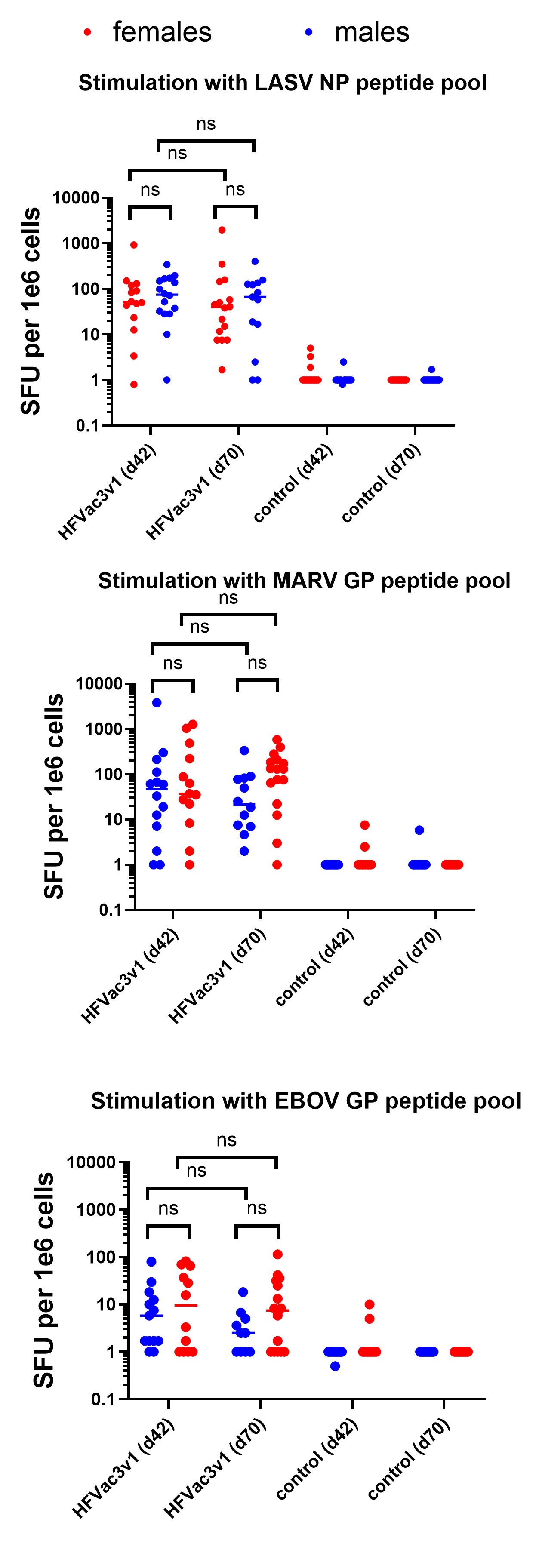
